# Comparative analysis based on shared amplicon sequence variants reveals that cohabitation influences gut microbiota sharing between humans and dogs

**DOI:** 10.1101/2024.04.11.589141

**Authors:** Yutaro Ito, Miho Nagasawa, Kahori Koyama, Kohei Ito, Takefumi Kikusui

**Author notes:** **Equal contributions:** These authors contributed equally to this work and share first authorship (Yutaro Ito, Miho Nagasawa).

## Abstract

**Introduction:** The One Health concept is a comprehensive understanding of the interaction between humans, animals, and the environment. The cohabitation of humans and dogs positively affects their physical, mental, and social well-being. It is recognized as an essential factor from the One Health perspective. Furthermore, a healthy balance in the gut microbiome is essential for good health, and the sharing of gut microbes between humans and dogs may positively impact the health of both hosts. Therefore, elucidating the sharing of gut bacteria between humans and dogs is important for understanding One Health. However, most studies have examined sharing at the taxonomic level, and it remains unclear whether the same bacteria are transferred between humans and dogs, and whether they mutually influence each other.

**Methods:** Here, microbiome analysis and 16S rRNA gene amplicon sequence variant (ASV) analysis was performed to address this question.

**Results:** 16S rRNA gene ASVs analysis indicated that gut microbes have been transferred between humans and dogs. The overall structure of the gut microbiota within human-dog pairs remained unchanged after three months of adaptation. However, some ASVs were shared within human-dog pairs. Many shared ASVs were highly abundant within each host, and this high abundance may be considered a factor that influences bacterial transfer between hosts.

**Discussion:** Our results demonstrate the possibility of the direct transfer of gut bacteria between humans and dogs, providing an important insight for achieving good health through human-dog cohabitation.

## 1 Introduction

The concept of “One Health” is based on the comprehensive understanding that humans, animals, and the environment surrounding them are interconnected. It is a cross-disciplinary approach to solving problems through collaboration among people involved in human, animal, and environmental health. Therefore, discussions are underway to promote the comprehensive health of pets, who spend considerable time in the same environment as humans, from the perspective of the One Health Triad (1,2). One of the central issues is the sharing of microbes and infectious diseases; issues related to zoonoses, in particular, have consistently received high attention.

Human residential microbiomes coexist in various body sites, such as the gut, skin, lungs, and oral cavity. It is estimated that the total number of bacteria in the 70 kg “reference man” is 38 trillion cells (3). The gut microbiome is the primary factor maintaining health. An imbalance due to external changes can lead to the development of cardiovascular diseases, cancer, respiratory diseases, diabetes, inflammatory bowel disease (IBD), brain diseases, chronic kidney diseases, and liver diseases (4,5).

Human residential bacteria are substantially affected by multiple factors in the external environment, including living spaces (6,7). Pets sharing living environments with humans have been reported to be a considerable factor influencing the taxonomic composition and phylogenetic diversity of the human gut and skin microbiomes via direct or indirect microbial transfer (8–14). Contact between humans and pets alters the composition of gut bacteria and potentially reduces the risk of allergic diseases in infants (9,15,16), and metabolic syndromes (17). The dog is regarded as the first domesticated animal (18). Domestic dogs are in daily contact with their owners and share their living environments. Regarding mental health, some studies have shown that dog ownership has an impact on improving human well-being through changes in physiological functions, such as endocrine regulation (19–21). Another study reported that the modification of dog microbiota by specific probiotics was reflected in the gut microbiome of children (22). Therefore, we must understand the impact of ecological interactions on microbial structures, and how their transfer across the One Health Triad affects human and dog health.

Although the effect of dogs on the human microbiome is considered substantial, most studies have discussed this at the taxonomic level. The direct transfer of gut microbes from dogs or coincidental sharing of the same taxa between humans and dogs is unclear. The dog gut microbiome was similar to the human gut microbiome, with 63% mapping to the human gene catalog (23), suggesting a possible interaction. In this study, we hypothesized that spending time with owners leads to microbial sharing between humans and dogs, resulting in similar gut microbiomes. To test this hypothesis, we analyzed microbial sharing at the amplicon sequence level.

## 2 Materials and methods

### 2.1 Study design

We examined 28 individual families with dogs and human participants between the ages of 20 and 72 years (48.5+/-15.7; 13 males and 15 females), and dogs between the ages of 1 and 10 years old (4.4+/- 2.6; 18 males and 10 females; 5 pure breed, 23 mix breed; 15 stray, 8 breeder, 3 house dog, 2 unknown). The dogs were obtained from shelters and breeders and were adopted to new families. Detailed characteristics of the study are summarized in Table 1. Fecal samples were collected from both humans and dogs. Fecal samples were collected from the dogs at the facility where they were kept for two to three months prior to adoption and from the owners one week prior to adoption, as well as from both the owners and dogs at two weeks, one month, and three months after adoption. The owners defecated using a fecal inspection sheet (Nagasale 0-9761-01, AS ONE Co. ltd., Osaka, Japan) placed in a toilet bowl. A portion of the fecal sample was scooped out without contact with water using disposable chopsticks, placed in a tube (CELL reactor filter cap centrifuge tube, 227245, Greiner Bio-one, Tokyo, Japan), and covered with a lid. The tube containing the fecal sample was placed in a pouch bag (A-58, Mitsubishi Chemical Corporation, Tokyo, Japan) together with AnaeroPack™-Anaero (A-03, Mitsubishi Gas Chemical Corporation, Tokyo, Japan) and made anaerobic. When dogs defecated, such as during a walk, an uncontaminated portion of soil or sand was collected using disposable chopsticks and placed in a tube under the same anaerobic conditions as those used for humans. After collection, the samples were placed in a cooler box with frozen refrigerant, sealed, and refrigerated until the following day. Samples from three of the 28 pairs were stored in preservation solutions (RNAlater™ Stabilization Solution, AM7022, Invitrogen, Thermo Fisher Scientific Inc.) due to changes in transportation methods. This method is comparable to the immediate freezing (24). Immediately after defecation, a small amount of feces was removed with a disposable microspatula (1-9404-02, AS ONE Co. Ltd., Osaka, Japan), placed in a 1.5 mL tube containing RNAlater and sealed. These samples are stored in a freezer at -80°C in the laboratory until analysis.

**Table 1.**
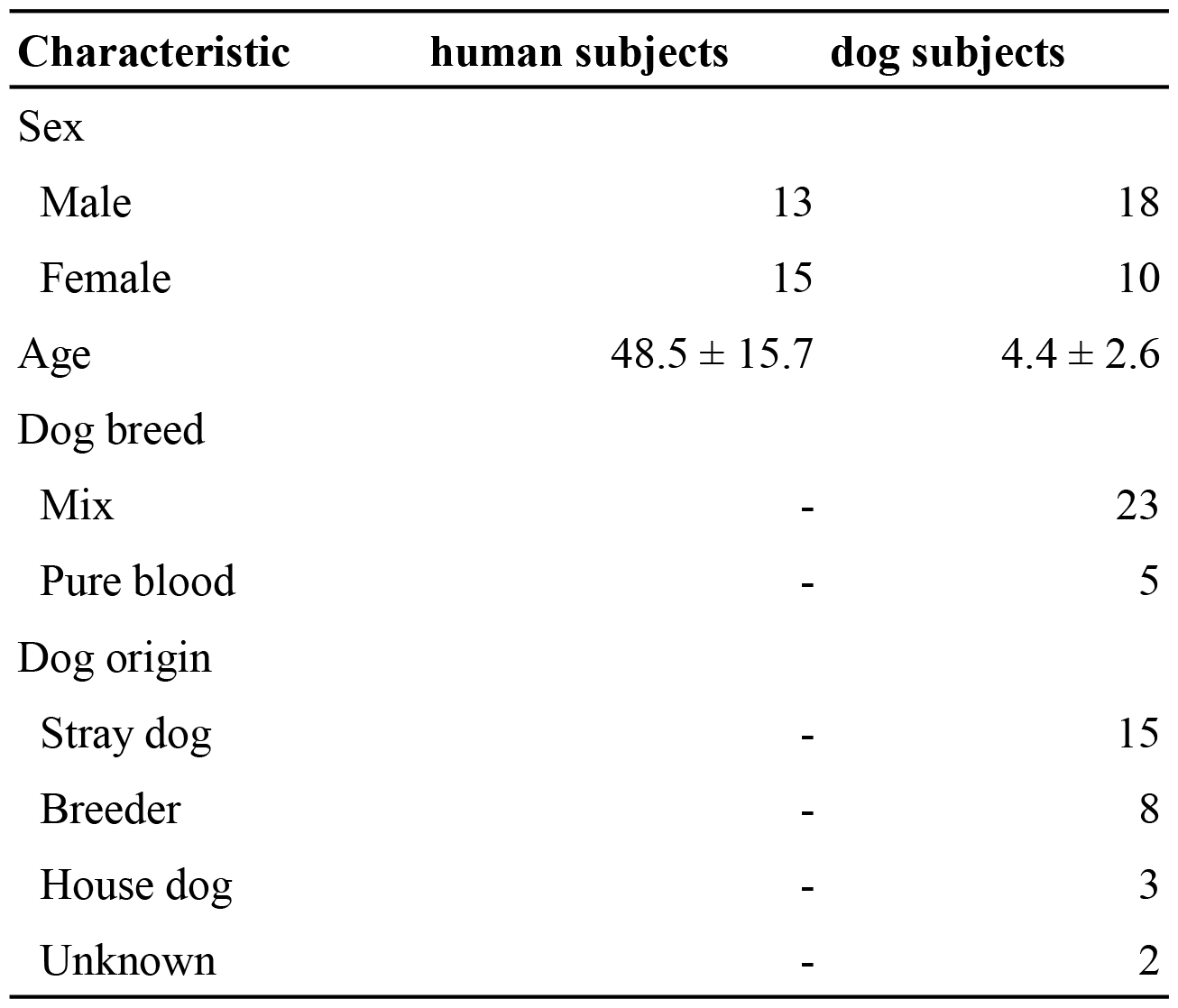
Characteristics of human and dog subjects.

### 2.2 Ethics

The study protocol was approved by the Animal Ethics Committee of Azabu University (#210325-12) and the Ethical Committee for Medical and Health Research Involving Human Subjects of Azabu University (#097). All procedures were conducted in accordance with the guidelines and regulations of the Ethics Committee. Informed consent was obtained from all participants, who were provided with detailed information about the study’s objectives, procedures, potential risks, and their right to withdraw at any time without penalty. To protect the privacy of participants, all personal identifiers were removed, and data were coded to maintain confidentiality and anonymity.

### 2.3 Total DNA extraction and high-throughput sequencing

Samples were treated with Lysis Solution F (NIPPON GENE Co., Ltd., Tokyo, Japan) and homogenized for 2 min at 1,500 rpm using a Shake Master Neo (Biomedical Science, Japan). The suspension was heat-treated at 65°C for 10 min, and centrifuged for 2 min at 12,000 × g. DNA was extracted from the separated supernatant using a Lab-Aid824s DNA Extraction Kit (Zeesan Biotech Co., China) according to the manufacturer’s protocol. In addition, PCR reactions were conducted with the bacterial universal primers 1st-341f_MIX (5′-ACACTCTTTCCCTACACGACGCTCTTCCGATCT-NNNNN-CCTACGGGNGGCWGCAG-3′) and 1st-805r_MIX (5′-GTGACTGGAGTTCAGACGTGTGCTCTTCCGATCT-NNNNN-GACTACHVGGGTATCTAATCC-3′), to amplify the V3-V4 of the 16S rRNA gene. The thermal conditions were 94°C for 2 min, followed by 98°C for 10 s, 55°C for 30 s, and 68°C for 30 s, with a final extension at 68°C for 7 min. DNA samples, library preparation, and amplicon sequencing were performed using 300-bp paired-end sequencing on the MiSeq Reagent Kit v3 (Illumina Inc., San Diego, CA, USA) and Illumina MiSeq platform (Illumina Inc., San Diego, CA, USA.) at the Bioengineering Lab. Co., Ltd. (Kanagawa, Japan).

### 2.4 Microbiome analysis

Microbiome analysis was performed as previously described (25). Briefly, the raw FASTQ data were imported into the QIIME2 platform (version 2023.5) as qza files (26). Denoising sequences and quality control were performed using QIIME dada2, and sequences were converted into amplicon sequence variants (ASVs) (27). ASVs were assigned to the SILVA database SSU 138.1 using the QIIME feature-classifier classification scikit-learn package (28,29). Subsequent analyses excluded ASVs classified as mitochondrial, chloroplast, or unassigned. To evaluate the effect of sequence read counts on microbiome diversity, we plotted changes in the Shannon diversity index over a range of read counts from 0 to 10,000 using rarefaction curves. In the rarefaction curves, the number of ASVs leveled off when the number of reads reached approximately 4,000 (Supplementary Figure 1). Beta diversity indices weighted by UniFrac distances were calculated, and the microbial community structure differences between groups were visualized using principal coordinate analysis (PCoA). Data were visualized using ggplot2 (version 3.4.4) (30) and ggprism (version 1.0.4) (https://csdaw.github.io/ggprism/) (Creators Charlotte Dawson1 Show affiliations 1. University of Cambridge, no date; Wickham, no date) software.

### 2.5 Calculation of shared ASVs

Shared ASV analyses were performed as described previously (31). In this study, we defined shared ASVs as those shared by > 1.0% of the samples. The calculation was conducted using custom Python code (q2-shared_asv, https://github.com/biota-inc/q2-shared_asv) with 0.01 for --p-percentage.

### 2.6 Statistical analysis

Mann–Whitney U tests were used to compare alpha diversity (Shannon diversity index) and pairwise UniFrac distances. To compare differences in beta diversity (Weighted UniFrac distance) between samples, for all PERMANOVA analyses, 5000 trials were performed to assess statistical significance. All multiple testing corrections were performed by computing False Discovery Rate using the Benjamini–Hochberg method, and *Q*-values (adjusted *P*-values) < 0.05 were considered statistically significant. Statistical tests were performed using SciPy (version 1.9.3) (32) and Scikit-bio (version 0.5.9) (http://scikit-bio.org). To validate the abundance of genera in humans and dogs, we used the analysis of composition of microbiomes (ANCOM) (33).

## 3 Results

### 3.1 The taxonomic composition of gut microbiomes in humans and dogs

We analyzed the composition of the gut microbiota in humans and dogs. The most abundant genera in the human gut were *Bifidobacterium, Blautia, Streptococcus, Bacteroides*, and *Faecalibacterium* (Figure 1A). *Fusicatenibacter* was significantly abundant only in humans, as determined by ANCOM (Supplementary Table S1). The most abundant genera in dog gut were *Streptococcus, Blautia, Peptoclostridium, Fusobacterium*, and *Ruminococcus gnavus* (Figure 1B). *Peptocrostridium* and *Blautia* were significantly more abundant in dogs, compared to that in humans, using ANCOM (Supplementary Table S1). *Blautia* and *Streptococcus* were abundant in humans and dogs. The top five dominant genera in each host collectively represented 51.6% (Interquartile range (IQR) 42.0–63.7) in the human gut and 46.2% (IQR 33.0–63.7) in the dog gut, based on the median relative abundance (Supplementary Figure S1A). The Shannon diversity index, the most commonly used index to measure the alpha diversity of the gut microbiome (34), did not change throughout the three-month cohabitation period (Supplementary Figure S1B).

**Figure 1.**
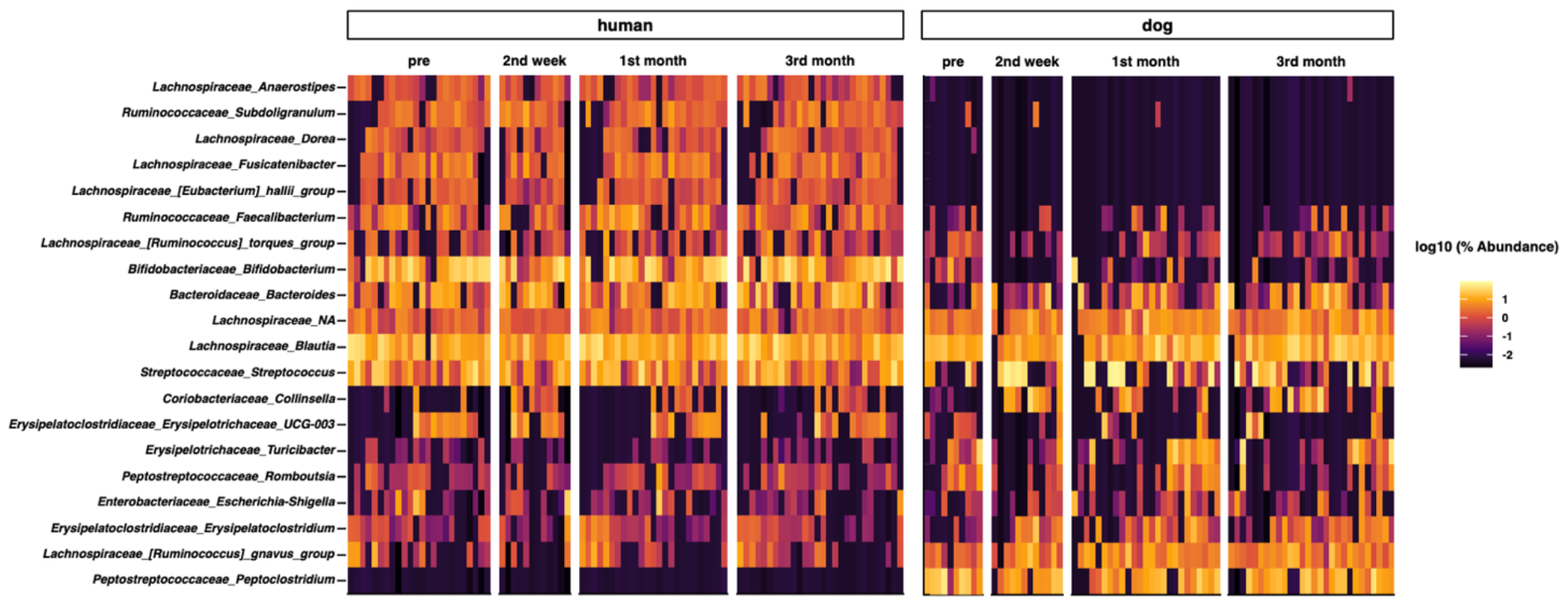
Taxon abundance heatmap at the genus level. Heatmap depicts the relative abundance (log10 scale) of the top 20 genera in humans and dogs.

### 3.2 The changes in microbial diversity and structures through the cohabitation of humans and dogs

We analyzed beta diversity to investigate the influence of cohabitation between humans and dogs on bacterial communities. The host species influenced the overall structure of the gut microbiota (*P* = 0.00020 based on PERMANOVA), as indicated by PCoA using the weighted UniFrac distance, whereas the duration of cohabitation did not have a notable impact (Figure 2A). The cohabitation period also did not affect the gut microbiota structure within human-dog pairs (Figure 2B). The weighted UniFrac distance was compared between the pre- and first month of shared living, and first month and third month of cohabitation (Figure 2C). Although the human gut microbiome had no discernible impact after cohabitation, it underwent significant alterations during the first month of cohabitation.

**Figure 2.**
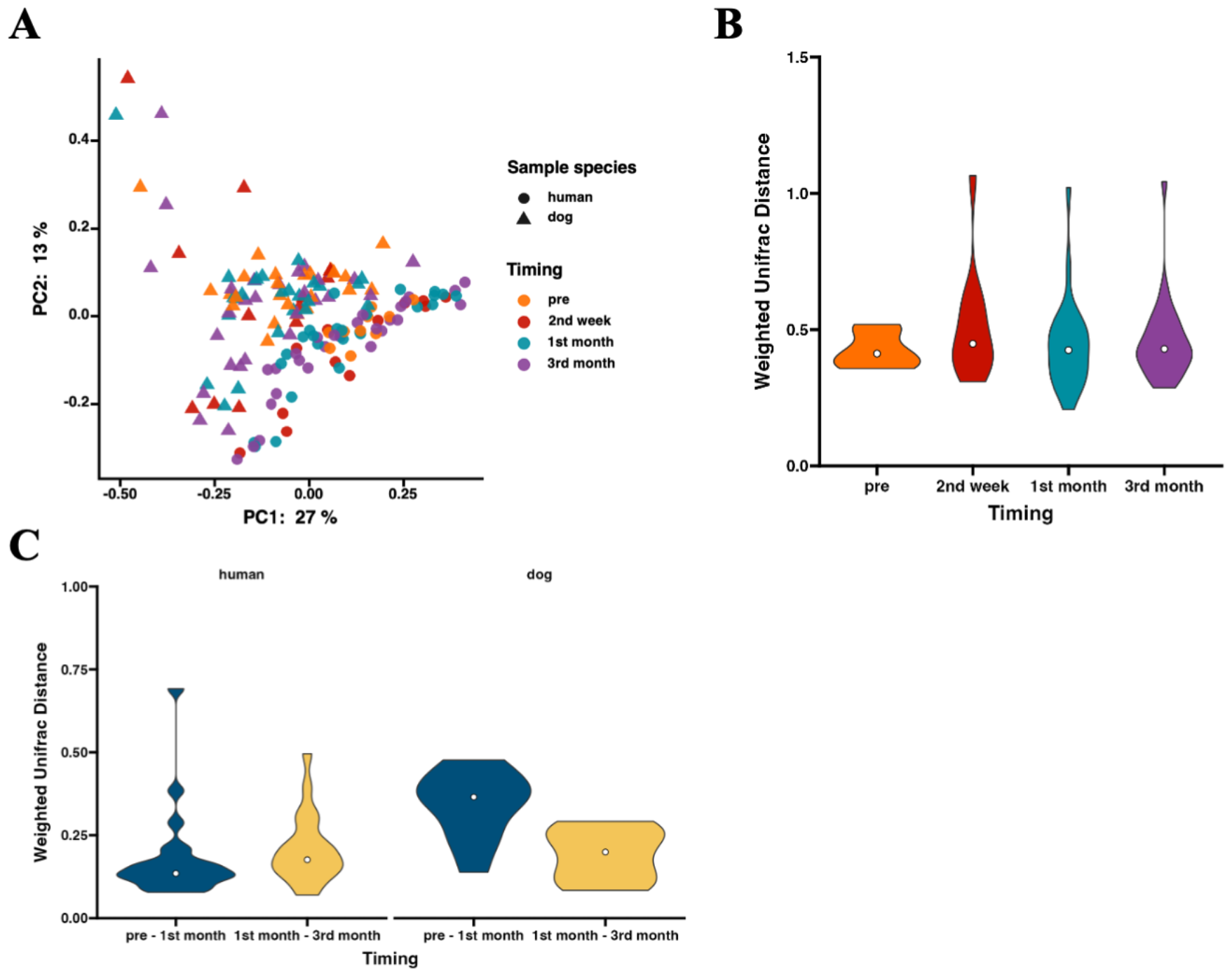
The gut microbiota structures in humans and dogs during each cohabitation period. Microbial profiles of gut microbiomes within human-dog pairs. (A) Principal coordinate analysis (PCoA) plot of human and dog gut microbiomes at each time point based on weighted UniFrac distance. (B) Violin plots of weighted UniFrac distance within human-dog pairs at pretest (human: n = 24, dog: n = 10), 2nd weeks (n = 12), 1st month (n = 25), and 3rd months (n = 28). Significance was calculated using the Kruskal-Wallis test. (C) Violin plots of weighted UniFrac distance between pre- and 1st month (n = 10) and between 1st month and 3rd months (n = 25) within human-dog pairs. Significance was calculated using the Mann-Whitney U test.

### 3.3 Time-series changes in shared ASVs of the gut microbiota between humans and dogs

Although the overall gut microbiomes within human-dog pairs were not influenced by shared living conditions, the possibility of sharing the same taxon at the ASV level within each pair was considered. A total of 5,709 ASVs were obtained from all samples. Shared ASV analysis revealed that only 11 ASVs were shared within human-dog pairs during the first and third months of cohabitation, but not the second week of cohabitation (Table 1). ASV001 and ASV002 were assigned to the *R. gnavas* group, the major bacterial genus in dog guts, and were shared across multiple pairs (Figure 3A). In one pair, these ASVs were exclusively identified in a dog sample at the first month and were later shared between humans and dogs at the third month. The other pairs shared ASVs at the same time point. ASV007, assigned to *Faecalibacterium*, was shared during the first month (Figure 3B). ASV was not detected at any other time points in either host strain. ASV010, assigned to *Streptococcus*, was detected only in the third month, with a relative abundance of 56.8% in the dog sample and was shared at this time point (Figure 3C). ASV005, which was assigned to *Blautia*, appeared simultaneously and was shared during the first month (Figure 3D).

**Figure 3.**
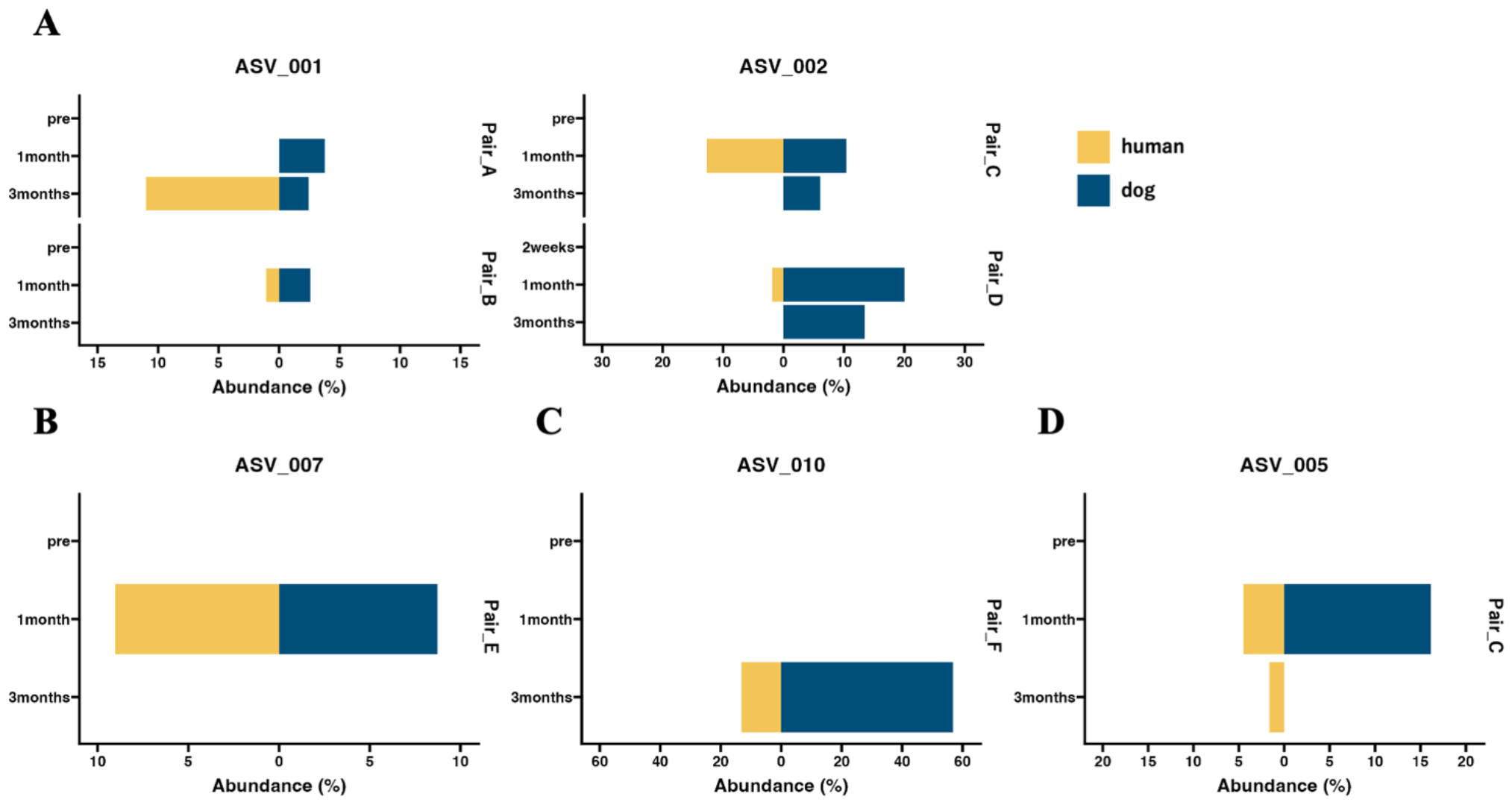
The abundance of shared ASVs within human-dog pairs at each time point. Butterfly chart of shared ASVs within human-dog pairs based on relative abundance. The pair IDs represent pairs of individuals, humans, and dogs sharing ASVs. (A) ASV_001 and ASV_002 were assigned to the *Ruminococcus gnavas* group. (B) ASV_007 was assigned to *Faecalibacterium*. (C) ASV_010, which was assigned to *Streptococcus*. (D) ASV_005 is assigned to *Blautia*.

## 4 Discussion

In the present study, we showed that the five dominant genera in each host collectively constituted approximately half of the relative abundance in their respective hosts. The top five dominant genera in the human gut microbiome, *Bifidobacterium, Blautia, Streptococcus, Bacteroides*, and *Faecalibacterium* have been reported as major components of the Japanese human gut microbiome (35). The genera abundant in the gut microbiome of dogs, including *Streptococcus, Blautia, Peptoclostridium*, and *Fusobacterium*, have also been identified as major components of the dog gut microbiome (36). *R. gnavas* has been reported as the most abundant species in the dog gut (23), which was also identified in this study. The human and dog gut microbiota in this study were considered to have no significant differences compared to those in previous reports.

We evaluated the Shannon diversity index to investigate the impact of human-dog cohabitation on community diversity. However, no variation was observed in cohabitation duration. Previous studies have reported that alpha diversities in the human gut microbiome do not show substantial differences (10,37), consistent with the current results. Beta diversity based on weighted UniFrac distances was compared over time, revealing no substantial changes in the overall microbial structures between pairs within the three-month cohabitation period. Similar to previous studies, in our study, the hosts (humans or dogs) were the main factors in explaining gut microbiota differences, and cohabitation did not seem to be one of the main factors affecting the overall gut microbiome composition (12,22). Finally, we analyzed the temporal variations in beta diversity to elucidate the changes in within-species microbial structures due to cohabitation. The weighted UniFrac distance of dog gut microbiomes between the pretest period and first month was significantly different from the distance between the first and third months. Changes in living conditions, such as diet and residence, when rescued dogs begin living with humans result in significant alterations in their gut microbiota during the early stages of cohabitation (38,39). However, no temporal changes were observed in the human gut microbiota due to cohabitation, suggesting that compared to dogs, there is a limited impact of environmental changes caused by cohabitation.

Although a correlation within human-dog pairs was not observed when considering the entire microbiome, we considered the possibility of shared individual bacteria between humans and dogs. Previous studies have compared changes in the gut microbiota due to cohabitation at the level of bacterial genera or OTUs (10,22,37). However, these analyses did not determine whether the same bacteria were transferred or shared. To precisely evaluate the sharing of the gut microbiota between humans and dogs, we conducted a shared ASV analysis, which is an approach for inferring the sharing ratio of the microbiome at the ASV level between samples and has been used previously (31,40,41). While the sharing of ASVs was not detectable after two weeks of cohabitation, it was observed in the first and third months. These results suggest that a cohabitation period of at least one month may be important for microbial sharing.

Among the 11 ASVs, six were classified as the dominant bacterial genera in the top five in each host. ASV001 and ASV002, assigned to the *R. gnavas* group, the common bacterial genera in the dog gut, were present in the dog samples of the shared pairs at more time points and were abundantly detected in the dog samples (Supplementary Table 2). These results suggested that the two ASVs were transferred from dogs to humans. Previous studies have demonstrated that *Ruminococcus* group 2 is more abundant in the guts of children with dogs, suggesting that *Ruminococcus* is easily transferred from dogs to humans (22). The *R. gnavus* group, found in abundance in the feces of human patients with IBD, produces polysaccharides and triggers the secretion of TNF-α from dendritic cells (42). The transfer of the *R. gnavus* group from dogs to humans is speculated to negatively affect human health. ASV007 assigned to *Faecalibacterium*, the major bacterial genus in the human gut, was shared in one pair and mainly detected in human samples (Supplementary Table S2). These results imply that ASV are shared between humans and dogs. In the human gut, *Faecalibacterium* is a beneficial bacterium, and may have beneficial effects when transferred to dogs (43). ASV010, assigned to *Streptococcus* mainly existed in dog samples (Supplementary Table S2) and was detected in the third month, with a relative abundance of 56.8% in the dog sample. It is considered that ASV010 became a major component of the dog gut microbiome due to factors such as chronic inflammatory enteropathy (44) and is subsequently transmitted to humans. ASV005, assigned to *Blautia*, was significantly detected in the dog gut based on ANCOM, while *Blautia* is a genus abundantly present in both hosts. These results suggested that this ASV is shared between dogs and humans. Notably, this ASV was detected exclusively in the human gut two months after sharing. *Blautia* is the second most abundant genus in humans, and it is possible that *Blautia* transferred from dogs to readily colonize humans. OTUs classified as *Blautia* become more abundant in the human gut due to cohabitation with pets, supporting this hypothesis (37). *Blautia* is recognized for its potential probiotic function in the human gut and it is speculated that transferring *Blautia* from dogs to humans may have beneficial effects (45). In conclusion, the pattern of this shared ASV suggests that the mutual sharing of bacteria between humans and dogs and high abundance are crucial factors in interhost microbial transfer.

**Table 2.** Genera of shared ASVs and the number of pairs sharing ASVs within humans-dog at each time point.

| ASV ID | Assign taxonomy | 2nd week | The number of shared pairs |  |
| --- | --- | --- | --- | --- |
|  |  |  | 1st month | 3rd month |
| ASV_001 | *Ruminococcus gnavus* group | - | 1 | 1 |
| ASV_002 | *Ruminococcus gnavus* group | - | 2 | - |
| ASV_003 | *Prevotella_9* | - | 1 | - |
| ASV_004 | *Prevotella_9* | - | 1 | - |
| ASV_005 | *Blautia* | - | 1 | - |
| ASV_006 | *Erysipelactoclostridium* | - | - | 1 |
| ASV_007 | *Facealibacterium* | - | 1 | - |
| ASV_008 | *Fusobacterium* | - | - | 1 |
| ASV_009 | Unknown (*Lachnospiraseae*) | - | - | 1 |
| ASV_010 | *Streptococcus* | - | - | 1 |
| ASV_011 | *Suttera* | - | - | 1 |

In this study, we aimed to analyze the interactions between the gut microbiota of adult humans and dogs. Consequently, there were very few (only 11) shared ASVs. Infants exposed to dogs at an early age have altered gut microbiota, which supports a potential mechanism explaining reduced atopy and asthma risk (46). The effect of microbial transfer may depend on the host age. Additionally, dog ownership increases the similarity of the skin microbiota between humans and dogs, rather than the gut microbiota (12). The closed nature of the intestinal environment may reduce the probability of bacterial transfer between hosts. In the future, it may be important to evaluate shared ASVs in various age groups and locations to better understand the interactions between humans and dogs.

This study has three limitations. First, we used amplicon sequences of the V3-V4 region of the 16S rRNA gene, which limited bacteria identification to the genus level (29). PCR amplification bias and differences in DNA extraction methods affect the accuracy of the relative composition of the gut microbiome (47,48). Shared ASV analysis is a convenient way to track microbial sources, and the full-length 16S rRNA gene can be used to predict microbial sharing between samples more rigorously. The lineages of bacterial strains between the two groups can be compared using metagenome-assembled genomes, which show more precision in tracking at the whole-genome level, not just the restricted specific hypervariable regions of the 16S rRNA gene. The second limitation is an imbalance in the sample size at each time point. Unlike the first and third month, where 25 and 28 samples were collected, respectively, we conducted the analysis using only 10 samples collected during the second week. It cannot be ruled out that the lack of shared ASVs in the second week samples may be attributed to the small sample size. Finally, we still need to demonstrate the impact of shared ASVs on each host. The shared ASVs analysis revealed the possibility of bacterial transfer. The influence of bacterial sharing on the health of each host could not be clarified. We believe that additional experiments, such as animal experiments using isolated bacteria demonstrated in this study, are necessary.

In conclusion, this study combined 16S rRNA gene amplicon and shared ASV analyses to provide high-resolution evidence of gut microbiome transfer during cohabitation between humans and dogs. ASVs shared in the gut exhibited a high relative abundance in each host, suggesting that ASV sharing is more likely to occur in the dominant taxon. Many ASVs that were confirmed to be shared were the dominant taxa in each host. A larger sample size is needed in future studies to differentiate the effects of different living environments, dog breeds, host sex, host age, and time spent with dogs. Further analysis is required to determine the relevance of ASVs specifically shared by each individual to subsequent health risks from the perspective of One Health.

## 6 Author contributions

All the authors conceived the study, contributed to the manuscript, and approved the submitted version.

Yutaro Ito: drafted the original manuscript and performed microbiome analysis.

Miho Nagasawa: handled research design and planning.

Kahori Koyama: conducted data collection.

Kohei Ito: drafted the original manuscript and performed microbiome analysis.

Takefumi Kikusui: provided supervision of research.

## 8 Acknowledgments

All authors thank Morgenrot Inc. for providing the computational environment for the analysis. We would like to thank Editage (www.editage.jp) for English language editing.

## 10 Data availability statement

The datasets generated through 16S rRNA amplicon sequencing are available and deposited in the NCBI Sequence Read Archive (SRA) database under accession numbers DRR542122-DRR542285 and BioProject PRJDB17718.

