## Supplemental materials for "Comparative analysis based on shared amplicon sequence variants reveals that cohabitation influences gut microbiota sharing between humans and dogs"

### *Supplementary Material*

#### 1. Supplementary Figures

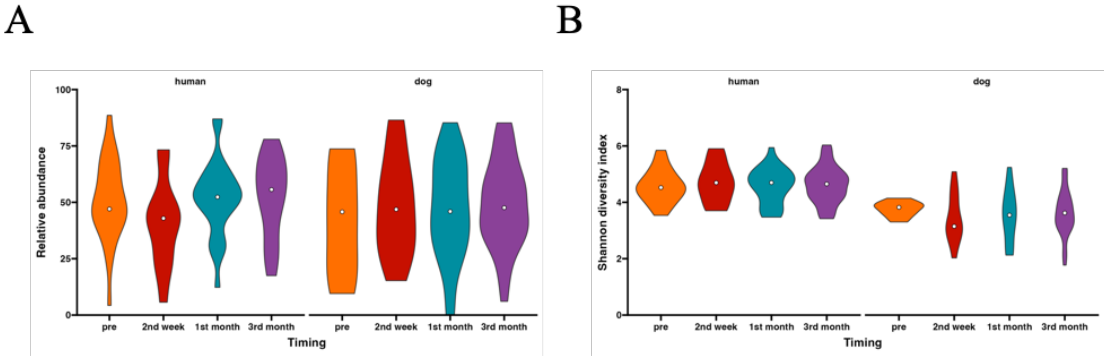


**Supplementary Figure 1.** The composition and diversity of intestinal microbes in humans and dogs. The composition and diversity of intestinal microbes in humans and dogs before and after cohabitation. (A) Violin plots of total relative abundance of the top five dominant genera in each host at pretest (human: n = 24, dog: n = 10), second week (n = 12), first month (n = 25), and third month (n = 28). (B) The Shannon diversity index in each host at each specified time point is indicated.

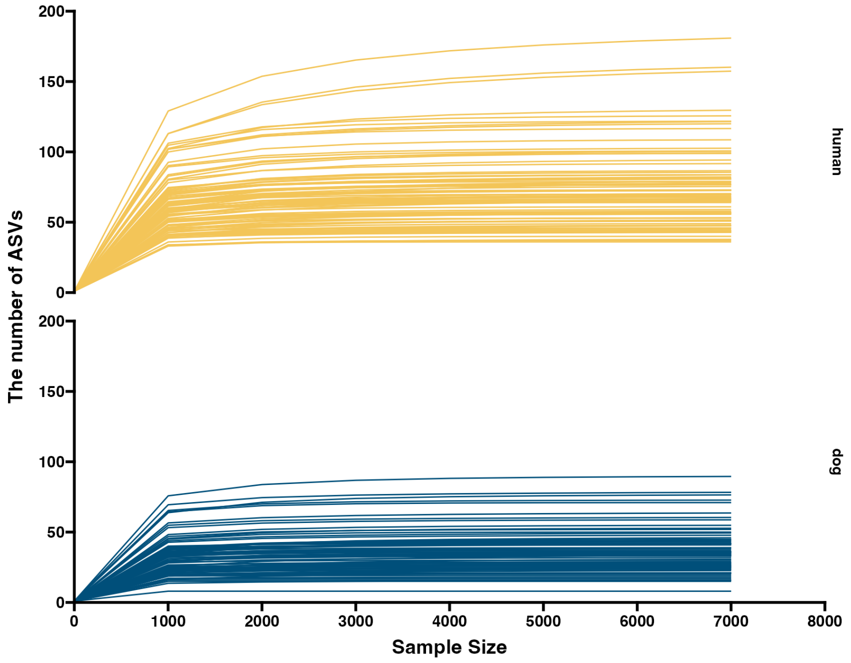


**Supplementary Figure 2. Rarefaction curves for the number of observed ASVs by sample size.** Rarefaction curves for the human and dog fecal samples. The blue line represents the rarefaction curve for the human gut microbiome, while the red line corresponds to the rarefaction curve for the dog gut microbiome. Each curve illustrates the observed number of ASVs at varying sequencing sample sizes. ASV, amplicon sequence variant

#### 2. Supplementary Tables

**Supplementary Table 1.** Statistically significant ANCOM results at the genus level. Relative abundance across all samples or ASVs within a group was summed.

| ASV ID | Taxonomy | clr | W-statistic value | Number of features (100 percentile) human | Number of features (100 percentile) dog |
|---|---|---|---|---|---|
| 85ed0d0f579237bd0a2f8 b98c28b3a5c | *Peptoclostridium* | 2.884 | 366 | 1 | 3803 |
| 27e2aa13ac8d9a01ddf28 63741c5cbcb | *Blautia* | 2.672 | 360 | 1 | 5677 |
| fa18371d7062793ea19a3 15790b69117 | *Peptoclostridium* | 2.494 | 360 | 641 | 5657 |
| 8e7b50632385bb16ea32e 2ae7f5c1a50 | *Fusicatenibacter* | -1.949 | 351 | 2067 | 1 |
| 1c911e95f53361d9972bb d73b40ea416 | *Blautia* | 1.598 | 331 | 449 | 1670 |
| fb7ae9e4b7fb258deaba2e 96fe851d32 | *Lachnospiraceae sp.* | 1.675 | 331 | 1 | 1725 |

**ANCOM, analysis of composition of microbiomes; ASV, amplicon sequence variant**

**Supplementary Table 2.** The ratio of samples in which ASVs with a detection rate of > 1% were confirmed in humans and dogs.

| ASV ID | human samples | dog samples |
| --- | --- | --- |
| ASV_001 | 0.079 | 0.31 |
| ASV_002 | 0.079 | 0.29 |
| ASV_003 | 0.045 | 0.013 |
| ASV_004 | 0.045 | 0.013 |
| ASV_005 | 0.023 | 0.387 |
| ASV_006 | 0.023 | 0.027 |
| ASV_007 | 0.045 | 0.027 |
| ASV_008 | 0.023 | 0.053 |
| ASV_009 | 0.023 | 0.187 |
| ASV_010 | 0.034 | 0.147 |
| ASV_011 | 0.011 | 0.013 |
